# High-sensitivity assessment of phagocytosis by persistent association-based normalization

**DOI:** 10.1101/827568

**Authors:** Therese de Neergaard, Martin Sundwall, Pontus Nordenfelt

**Affiliations:** Division of Infection Medicine, Department of Clinical Sciences, Faculty of Medicine, Lund University, Lund, Sweden

**Keywords:** Phagocytosis, Quantitative, Bacteria, Flow cytometry

## Abstract

Phagocytosis is measured as a functional outcome in many research fields but can be challenging to quantify accurately, with no robust method available for cross-laboratory reproducibility. Here, we identified a simple, measurable parameter – persistent prey-phagocyte association – to use for normalization and dose-response analysis. We apply this in a straightforward analytical method, persistent association-based normalization (PAN), where first the multiplicity of prey (MOP) ratio needed to elicit half of the phagocytes to associate persistently (MOP_50_) is determined. MOP_50_ is then applied to normalize for experimental factors, with adhesion and internalization analyzed separately. We use THP-1 cells and different prey and opsonization conditions to compare the PAN method to standard ways of assessing phagocytosis, and find it better in all cases, with increased robustness, sensitivity, and reproducibility. The approach is easily incorporated into most existing phagocytosis assays and allows for reproducibly comparing results across experiments and laboratories with high sensitivity.

## Introduction

Phagocytosis is an ancient receptor-driven cellular process that involves the engulfment of a particular prey by a cell with phagocytic capability (Flannagan et al., 2012). In metazoans, it is essential for the immune system as well as for homeostasis by cleaning up cellular debris and waste (Lim et al., 2017). As it is such a ubiquitous process, it is natural that many research fields study or use phagocytosis as a functional readout, and ever since Metchnikoff discovered the phenomenon (Metschnikoff, 1884), researchers have come up with different ways to assess and quantify phagocytosis.

Phagocytosis has typically been measured either through direct observation with electron or light microscopy or through various indirect methods where the prey is labeled with a marker, such as radiation (Nordenfelt, 1970) or fluorescence (Hed, 1977). Several approaches are able to discriminate between prey that are on the outside or the inside of the phagocyte, typically by double labeling (Heesemann and Laufs, 1985; Nordenfelt et al., 2009), quenching (Hed et al., 1987)), or by fluorescent color change induced by the intracellular environment (Miksa et al., 2009). Arguably, the most common instrument used today is the flow cytometer, where fluorescently labeled prey and phagocytes can be quantified in a high-throughput manner (Ackerman et al., 2011), and sometimes in combination with fluorescence microscopy for qualitative confirmation (Dunn and Tyrer, 1981). Another common approach is the colony-forming unit-based assay (Lancefield, 1959; Todd, 1927), where phagocytosis is indirectly assessed through phagocyte-mediated killing, and typically employed when the question is more focused on the outcome of the phagocytic process, rather than the process itself.

Ideally, we would like to be able to draw general conclusions about phagocytosis and compare results across different biological systems. We know that phagocyte characteristics differ significantly, including for the two most common human ones, neutrophils and macrophages (Nordenfelt and Tapper, 2011). We also know that prey characteristics such as shape and size (Champion and Mitragotri, 2006), as well as surface properties including opsonization density (Zhang et al., 2010) and localization vary even more (Bakalar et al., 2018; Nordenfelt et al., 2012), and that the microenvironment plays a vital part in changing the prey characteristics (Lood et al., 2015). Besides the variability that biology will impart, there are also well-known physical and experimental factors that impact phagocytosis assays (Ponder, 1927), including but not limited to, temperature, time, volume, and the ratio of prey to phagocyte. Independent from method principle and regardless of the specific assay, there is currently no standardized way to reduce the impact of experimental factors. There is also no standard way to define a phagocytic index or how to describe phagocytosis assays. This lack of standards, including inconsistent terminology, leads to difficulty in comparing across experiments, experimental systems, and laboratories, ultimately affecting assay reproducibility and sensitivity. If phagocytosis could be assessed in a more reproducible manner and at higher sensitivity, besides the obvious benefits of higher quality data, it would also enable us to ask more detailed mechanistic questions revolving this ancient process.

We hypothesize that by determining the proportion of phagocytes that actively associate with prey, most of the factors that cause experimental variation can be normalized for, and in the process, increase the biological signal that can be measured (**Fig. 1A**). If we view the phagocytes and prey as particles that can react together, we can infer from collision theory that the critical event is a persistent collision event, where two particles interact in such a way that a reaction occurs, and they remain together (Manley and Mason, 1952; Ponder, 1927). Association between prey and phagocyte can then be described in terms of a *persistent association* event, and the level of the association will be related to collision frequency (**Fig. 1A**). If a similar level of association across various experimental conditions will result in a similar level of phagocytosis, this shows that normalization is possible for experimental variation as long as the degree of prey-phagocyte association can be determined.

**Figure 1.**
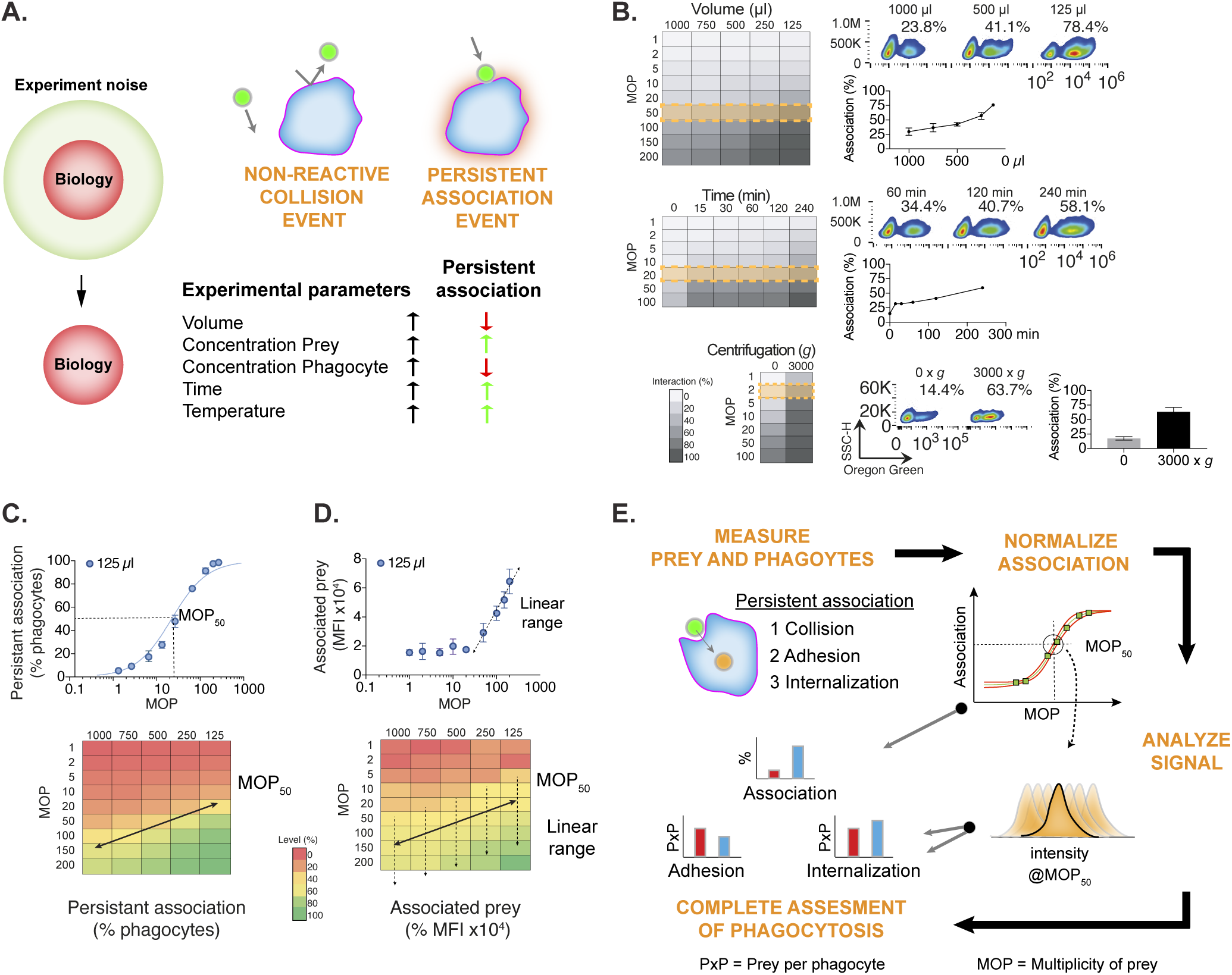
Persistent association frequency decides the relative outcome of phagocytic interaction. (A) To be able to enhance the biological signal in phagocytosis experiments, we wanted to define the most important parameters contributing to experimental noise. Based on collision theory we describe how experimental parameters affect the likelihood for finding a phagocyte interacting with a prey. When a prey-phagocyte collision occurs and interaction is persistent over the measurement time this is defined as a *persistent association* and can be quantified as the percentage of phagocytes associated with at least one prey. (B-D) THP-1 cells were incubated at 37°C with IgG-opsonized heat-killed *S. aureus* labeled with Oregon Green and measured with flow cytometry. (B) Heat maps of persistent association for a range of multiplicity of prey (MOP) in different experimental setups: volume (125-1000 µl, n=3), time (0-240 min, phagocytic volume 1500 µl, n=1) and centrifugation (0 and 3000 x *g*, n=3). Highlighting individual experiments with their corresponding flow cytometry scatter plots and quantified data. Data are represented as mean ± SD when possible. (C) The dynamic range of volume with respect to persistent association and MOP plotted in linear-logarithmic scale shows a sigmoidal dose-response relationship visualized with 125 µl and a heat map for each volume with the midpoint of the curve (MOP_50_) visualized in yellow. (D) The corresponding cell-associated prey expressed in mode fluorescence intensity (MFI) shows a linear relationship around MOP_50_. (E) A schematic overview outlining the proposed steps of normalized assessment of phagocytosis A suitable method that can measure phagocyte and prey signal is required, with discrimination of adhesion and internalization being optional. Phagocytosis is recorded over a range of MOPs, and then MOP50 is determined and used for quantification of normalized prey intensity. Depending on the labels used, intensity is then used to quantify associated, adhered and internalized prey, resulting in complete assessment of phagocytosis.

Here, we present a systematic approach to assess phagocytosis irrespective of the chosen experimental method. From collision theory, which concerns the likelihood that two particles collide and react, we can conclude that the phagocytic interaction step will have a considerable impact on experimental variation. We show that this can be normalized by employing dose-response curve analysis and comparing conditions at the same level of persistent prey-phagocyte association. Such normalization leads to increased assay robustness, reproducibility, and sensitivity. Persistent association normalization also allows for comparison across completely different experimental settings and laboratory conditions. Finally, we provide experimental tools and guidelines for different degrees of phagocytosis assessment, so that the researcher can match the method to the question being asked.

## Results

### Persistent association frequency decides the relative outcome of phagocytic interaction

Our aim was to establish a robust and standardized method to assess phagocytosis with high sensitivity. The method should also be able to discriminate between association, adhesion, and internalization. To achieve this, the method had to normalize for experimental factors modulating effective association – following a collision – between phagocyte and prey. If a phagocytosis experiment could be regarded as a reaction vessel with two reactants, collision theory describes the likelihood for a reaction to occur, based on collision frequency, and what parameters it depends on. For phagocytosis, we define such a reaction as a *persistent association*, with the critical parameters being reaction volume, reactant concentration, and reaction time (**Fig. 1A**). To have terminology that is relatable to phagocytosis, we introduce the term multiplicity of prey (MOP) to describe the ratio of particulate prey to phagocytes. The term MOP followed by a sub-index (MOP_X_) is to indicate the effective MOP to reach X percentage of persistent association, which is the result of an effective collision, and is similar to EC_50_ used in pharmacological studies (Holford and Sheiner, 1981). Finally, the term PxP is used to indicate number of prey per phagocyte. This results in the following equation to describe MOP_50_, which is a function of prey concentration, [MOP]: 

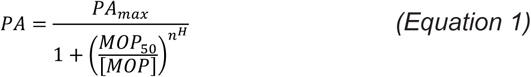

where PA is the observed persistent association, PA_max_ is the maximal PA at high prey concentrations, and where n^H^, the Hill coefficient (Hill, 1910), describes the slope of the curve.

To analyze how persistent association varies, we systematically tested the most important parameters derived from collision theory: volume, MOP, and time. To reduce the number of variables, we opted to use the THP-1 cell line for all experiments in this study, which are commonly used to measure phagocytosis (Chanput et al., 2014). They have a consistent expression of Fc receptors when kept at certain cell densities (<0.5 million cells/ml, (Ackerman et al., 2011)). As prey, we used Oregon Green-labeled *Staphylococcus aureus* opsonized with pooled polyclonal human IgG (IVIG). The results are summarized as heat maps in **Figure 1B** and show that several variables have potentially dramatic effects on the outcome. A few individual experiments are highlighted with their corresponding flow cytometry scatter plots and quantified data. The effect of changing volume is clear at MOP 50, where the persistent association increases from 30% to 70% when decreasing the volume. Similar for time, which is well-known (Nordenfelt, 2014), at MOP 20, increasing incubation time results in a doubling of the persistent association from 34% to 60%. Not surprisingly, the clearest effect can be found when phagocytes and preys prior to incubation are centrifuged together, at MOP 2, the interaction increases several-fold.

To evaluate how the dynamic range was affected by experimental factors we plotted volume with respect to MOP in linear-logarithmic (lin-log) dose-response curves and compared persistent association and corresponding cell-associated prey (**Fig. 1C-D**). The percentage of phagocytes that persistently associated with prey resulted in a sigmoidal relationship centered around MOP_50_ (**Fig. 1C**), and that this corresponded to a region with a linear relationship in terms of prey that associate with phagocytes (**Fig. 1D**). This trend could also clearly be seen when analyzing the whole range of MOPs and volumes, visualized as heat maps in **Figure 1C-D**. Since the dose-response relationship was consistent across such a large range of conditions, we postulated that interaction differences due to non-biological experimental factors, such as volume, could be normalized by using the characteristics of these types of curves.

In **Figure 1E** we summarize our proposed new way to assess phagocytosis, PAN (persistent association-based normalization) which works by normalizing interaction effects through estimating persistent prey-phagocyte association. In principle, a set number of phagocytes are incubated with a range of prey concentrations (MOPs). Data is acquired using any method that can distinguish between non-interacting and interacting phagocytes such as flow cytometry, imaging or other methods of choice. Then, a lin-log function is fitted to the data to estimate at what MOP half of the maximal persistent association is evoked (MOP_50_). Subsequently, the rest of the analyses are performed at MOP_50_ where persistent association-modulating experimental factors are normalized. This approach will allow for normalized assessment of phagocyte association with prey and can also include analysis of both adhesion and internalization frequencies.

### Experimental factors can be normalized by determining effective multiplicity of prey

To evaluate if we could use MOP_50_ to normalize experimental factors effect on collision frequency we designed an experiment where we could look at the change in persistent association separated from other factors. Given that the biology should be unaffected, we hypothesized that by changing experimental volume and nothing else, comparison at MOP_50_ should give the same number of prey per phagocyte (PxP), see **Figure 2A**.

**Figure 2.**
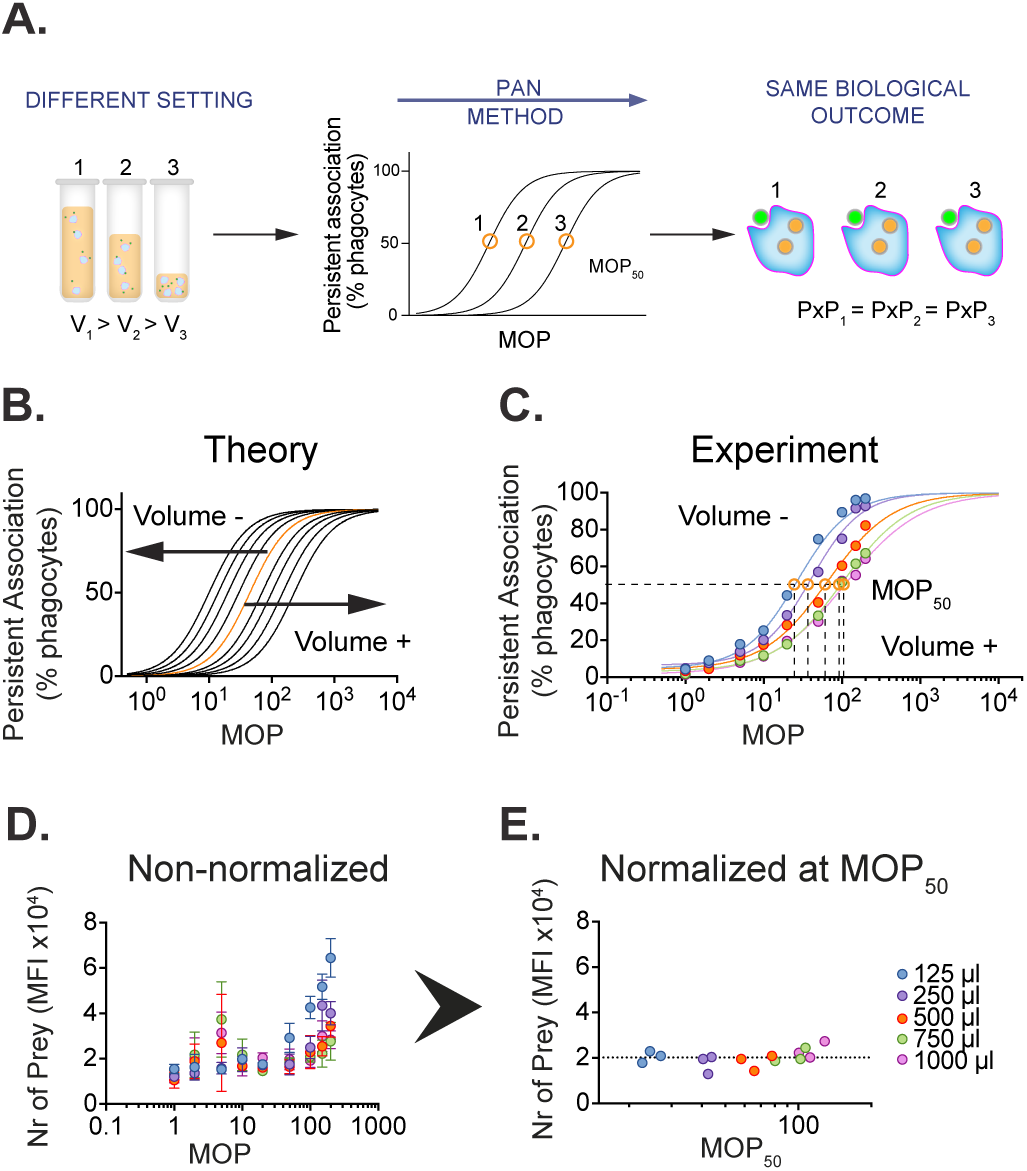
Experimental factors can be normalized by determining effective multiplicity of prey. (A) The schematic illustrates the hypothesis that if experimental factors, such as volume, can be normalized through MOP_50_ the same number of prey per phagocyte should be detected as biology should be consistent. (B) Theoretical persistent association curves for different volumes shows the curves shift to the right when volume is increased and to the left when it is decreased. (C-E) THP-1 cells were incubated with Oregon Green-labeled *S. aureus* for MOP 1-200 in volumes between 125-1000 µl, and measured with flow cytometry. (C) Representative dose-response curve of persistent association for each volume. The goodness of fit is R^2^ mean ± SD, n=3, 125 µl 0.99 ± 0.01, 250 µl 0.99 ± 0.01, 500 µl 0.99 ± 0.004, 750 µl 0.99 ± 0.01, 1000 µl 0.98 ± 0.01. (D-E) Number of prey per interacting phagocyte expressed in Mode Fluorescence Intensity (MFI) for each MOP (D, mean ± SD, n=3) and at MOP_50_ (E, each experiment visualized) normalizing the volume effect on persistent association.

**Figure 2B** shows the theoretical persistent association outcome of increasing (right shift) or decreasing (left shift) volume from the starting volume. The experimental data show similar results to the predicted outcome (**Fig. 2C**), with the association curves moving left or right, depending on the corresponding change in volume. The corresponding shifts in MOP_50_ is also indicated in the figure (**Fig. 2C**). The prey signal analysis shows a large range in observed prey signal per phagocyte, with up to 6-fold difference (**Fig. 2D**), which converges to very similar levels after applying MOP_50_-based normalization (**Fig. 2E**). Taken together, this suggests that experimental factors that cause a change in relative collision frequency can be normalized across experiments by estimating MOP_50_.

### Normalizing for persistent association frequency improves the sensitivity of phagocytosis assays

To address the question of whether normalizing for persistent association is an improvement for assay sensitivity and robustness, we wanted to compare the PAN-method to the current state-of-the-art. However, there are many ways to analyze phagocytosis and no single gold standard. We did a meta-analysis based on a random selection of 100 articles from 2013-2018 that included phagocytosis assessment of their studies (**Supp. Fig 1**). We found that we could group the methods into one indirect approach and three bona fide phagocytosis assessment levels (**Table 1, Fig. 3E**) and then proceeded to compare our method with them. The first level represents whole phagocytic population analysis, while the second also distinguishes between interacting and non-interacting phagocytes. The third, in addition to the second level, separate adhesion from the internalization of the prey. We place the approach described in this study as a fourth level, where the usage of MOP_50_ estimation to normalize for experimental factors can improve the detection of differences in association, adhesion, and internalization.

**Table 1.** Overview of different standard ways to assess phagocytosis. There is a wide range of different ways to assess phagocytosis and no single gold standard. Here we have grouped them based on their analysis principles into five different levels, we named them Indirect, A, B, C and D. Different types of indirect methods to study phagocytosis belongs to Level Indirect, such as plate-based killing assays. At Level A the interaction of the whole phagocytic population is evaluated, for example, an absorbance assay. With Level B non-interacting cells are separated from the interacting cells, and at Level C phagocytosis is further divided into adhesion and internalization. The PAN method, introduced here, requires that MOP_50_ is determined before quantifying phagocytosis, and is designated level D.

| Level | Description | Phagocytes | Type of data | Example |
| --- | --- | --- | --- | --- |
| Indirect | Imprecise | - | Unspecific | Killing assays |
| A | Generalizes | All | Population based | Absorbance assays |
| B | Separates | Associated | Prey vs no prey | Fluorescent prey |
| C | Specifies | Associated | Adhered vs internalized | Quenching |
| D | Normalizes | Associated | At MOP<sub>50</sub> | PAN-Method |

**Figure 3.**
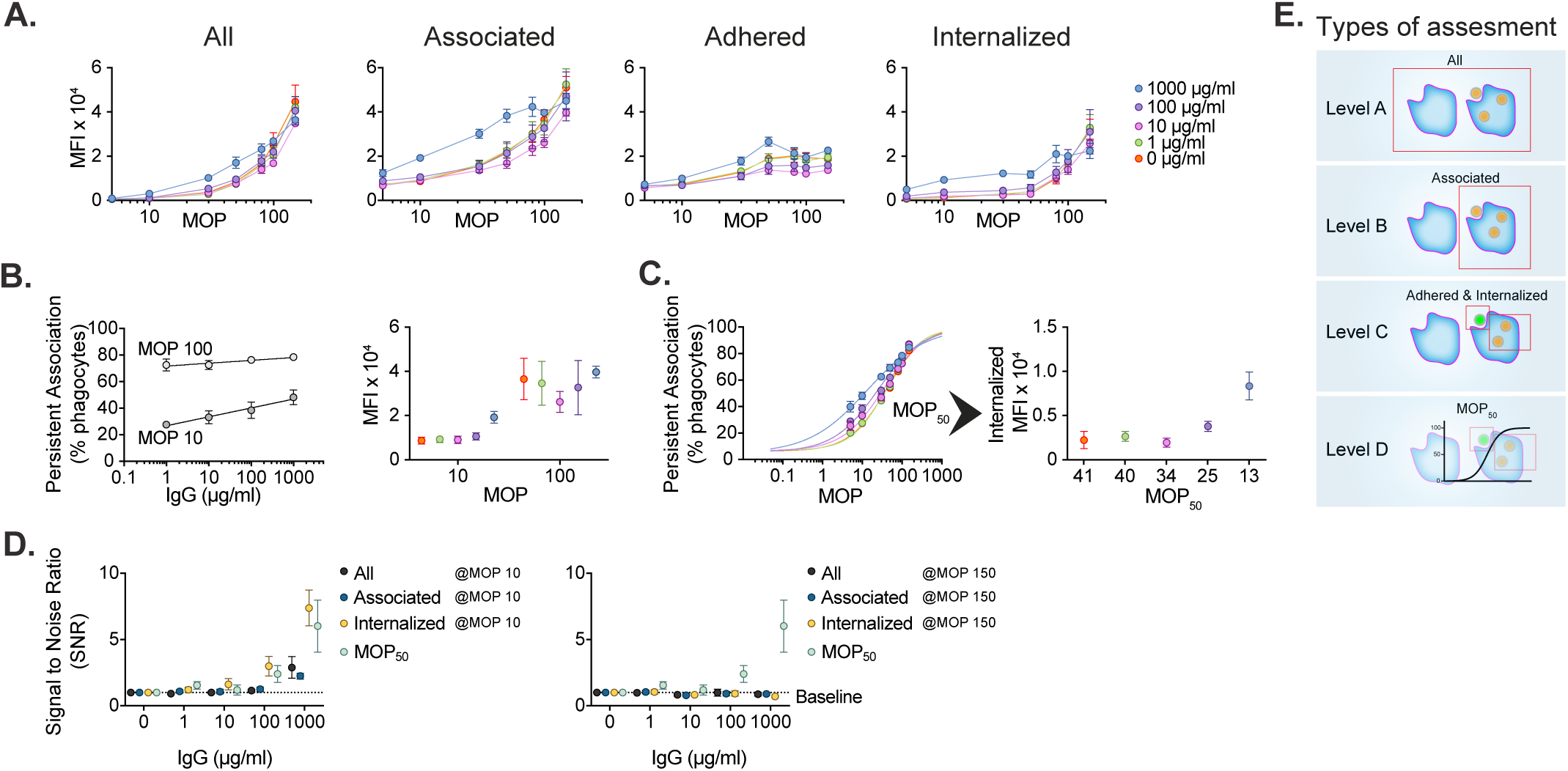
Normalizing for persistent association frequency improves the sensitivity of phagocytosis assays. (A-D) THP-1 cells were incubated (30 min, 150 µl, MOP 0-150, 37°C) with 1µm Far Red streptavidin beads opsonized with 0-1000 µg/ml of human polyclonal IgG. Alexa488-conjugated biotin was added post fixation to mark extracellular beads. Data were acquired through flow cytometry and are presented as mean ± SEM (A, C, D) ± SD (B), n=4. (A) The average number of prey per phagocyte expressed in Median Fluorescence Intensity (MFI). The internalized signal was quantified by subtracting the signal from the adhered beads from the associated beads signal. (B) Persistent association, the percentage of THP-1 cells interacting with beads, at two common MOP:s, 10 and 100 and the corresponding prey signal expressed in MFI. (C) The average persistent association curve for each IgG concentration (R^2^ mean ± SD, n=4: 0 µg/ml 0.99 ± 0.005, 1 µg/ml 0.99 ± 0.005, 10 µg/ml 0.98 ± 0.005, 100 µg/ml 0.99 ± 0.006, 1000 µg/ml 0.99 ± 0.003) and the average number of prey internalized per interacting phagocyte at MOP_50_. (D) Comparing the sensitivity of normalizing for persistent association with standard methods, illustrated in E, by quantifying the signal to noise ratio (SNR). SNR is defined as fold increase in prey signal compared to non-opsonized prey. This is compared at MOP 10 and 150 at standard ways to assess phagocytosis by different gating strategies; all phagocytes (black, Level A), associated phagocytes (blue, Level B), associated phagocytes with internalized prey signal (yellow, Level C) and internalized at MOP_50_ (green, Level D).

To compare these levels of phagocytosis assessment and evaluate whether MOP_50_-based normalization provides any benefits, we analyzed the same data set according to the different levels. Here, we kept the experimental parameters the same, and we varied a biological one, IgG density. Fluorescent streptavidin beads (far red) were opsonized with 0-10^3^ µg/ml IgG and after phagocytosis, the extracellular beads were coated with fluorescent Alexa 488-biotin (green). The data were acquired through flow cytometry where adhered beads were detected in both far-red and green channels, while internalized only in farred.

In **Figure 3A**, we show how the prey signal varies with IgG density, depending on which population of phagocytes is being analyzed. When all cells are included, there is almost no difference across the IgG densities, whereas more information is resolved when the analysis is focused on persistently associated phagocytes, as well as the further subdivision into adhered and internalized prey. This indicates that phagocytosis assessment gains sensitivity by increasing the level of analysis as described in **Table 1**.

We hypothesize that the biggest gain in sensitivity will come from MOP_50_ normalization. To exemplify the large effect MOP has on persistent association we compare two common MOPs used in the literature, 10 vs 100. At MOP 10, a clear difference in the association can be seen, and association increases with IgG density. This is nullified when the cells are overloaded at MOP 100 (**Fig. 3B**) resulting in the same range of persistent association for all IgG densities. Also, the resulting associated prey data are not consistent with expected results and varies depending on whether MOP 10 or MOP 100 is used. Consequently, if the analysis is performed at a fixed MOP independent of condition, which is most often the case in the literature, this will affect assay sensitivity. If the phagocyte association is instead normalized to a comparable MOP, the internalized prey signal increases (**Fig. 3C**). To quantify the sensitivity of different levels of analysis, we looked at signal to noise ratio (SNR) of the analysis, both at different MOPs as well as when using MOP_50_ normalization (**Fig. 3D**). We defined a biological SNR as a fold increase in prey association as compared to non-opsonized beads (0 µg/ml). This resulted in very low sensitivity for Level A and B analysis, with the only significant signal being detected at the highest degree of opsonization, and only at one selected MOP, which happened to be close to MOP_50_. Level C analysis worked well when the selected MOP was close to MOP_50_, but not at all when it was closer to MOP_90_. Additionally, MOP_50_-based analysis had a significantly higher SNR at all conditions, making it possible to detect a signal already at the lowest degree of opsonization (1 µg/ml). To summarize, the persistent association-normalized method represents a clear improvement compared to other standard methods in the field, with increased robustness and assay sensitivity.

### Association characteristics across opsonization density reveal biological differences

Determining association curves provides additional possibilities besides the increased robustness and sensitivity (**Fig. 4A**). The association curves can also give information about what to expect at low or high MOP situations, such as when few (MOP_10_) or many (MOP_90_) of the phagocytes are persistently associated with prey. Additionally, the saturation value of the curve (PA_max_), and the slope of the curve (Hill coefficient) is related to cooperative effects (Hill, 1910). This can reveal biological differences of phagocytosis between conditions, since physical parameters are being normalized for with the PAN-approach. Here, we apply these types of analyses to the simple scenario of changing the IgG density of the prey during phagocytosis.

**Figure 4.**
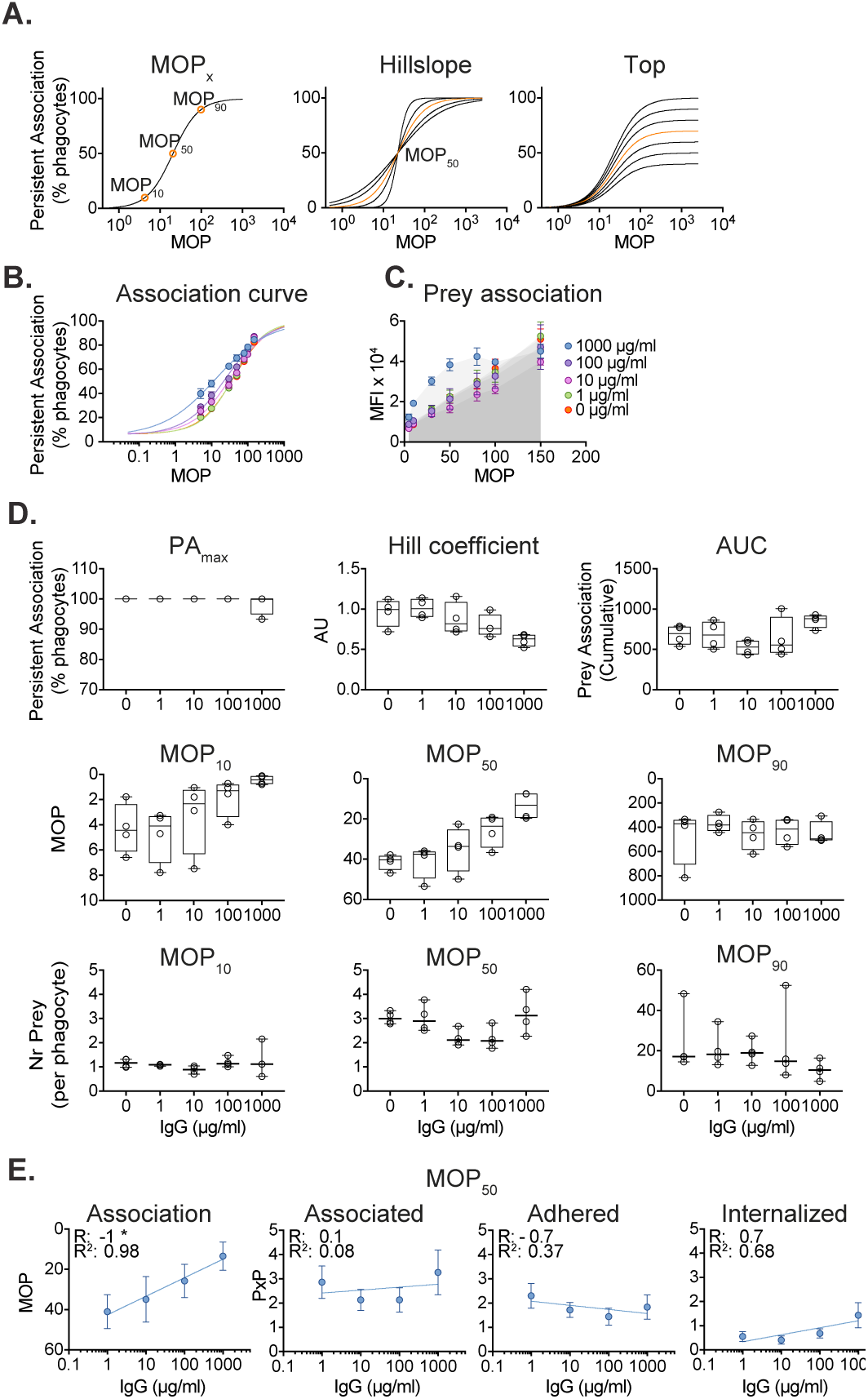
Association characteristics across opsonization density reveal biological differences. (A) Visualization of important characteristics of dose-response curves (Left) MOP_10_, MOP_50_, and MOP_90_ represent the MOP required to achieve corresponding degree of persistently associated phagocytes. (Middle) The curve shape is a measure of cooperative effects and is measured as Hill coefficient, with 1 being no cooperative effects, <1 negative cooperation and >1 positive cooperation. (Right) The top value of the curve indicates the maximum level of persistent association that the phagocytes can achieve. (B-E) Same experiment as Fig 3. Single beads were run separately allowing for conversion of the fluorescence signal to number of beads. Data were acquired through flow cytometry and are represented as mean ± SEM, n=4. (B) MOP_50_ was estimated through dose-response curves of persistent association (R^2^ mean ± SD, n=4: 0 µg/ml 0.99 ± 0.005, 1 µg/ml 0.99 ± 0.005, 10 µg/ml 0.98 ± 0.005, 100 µg/ml 0.99 ± 0.006, 1000 µg/ ml 0.99 ± 0.003) for each IgG density and (C) the corresponding prey MFI. (D) The characteristics of the dose-response curves for each IgG density as retrieved when using the template provided (Supp. Fig 4). All data points are shown, the distribution is visualized with box plots with whiskers ranging from minimum to maximum and a line marking the median, n=4. The last row shows all data points, their median with 95 % confidence interval, n=4 except at MOP_10_ where n=3 for 1000 µg/ml. (E) IgG density effect on phagocytosis, mean ± SD. The relationship was analyzed based on the mean with Spearman correlation as R with significant * p<0.05 and non-linear regression as R^2^. First the relationship of IgG and the MOP needed to reach MOP_50_ is presented. Followed by the number of beads associating, adhered and internalized per interacting phagocyte (PxP) at MOP_50_. Internalization was quantified by subtracting the adhered beads from the interacting ones.

The relationship between opsonin density and Fc receptor-based internalization is not completely clear with conflicting results in the literature (Gallo et al., 2010; Pacheco et al., 2013; Zhang et al., 2010), and the effect on persistent association has not been reported at all. **Figure 4B, 4C**, and **Supplementary Figure 2C** shows persistent association curves, prey association with phagocytes and examples of flow cytometry scatter plots, respectively, at different MOPs. This data shows an expected increase in association for both prey and phagocytes, which we analyze more carefully in **Figure 4D** based on curve parameters. The top row indicates that the maximum association value is barely affected by IgG density, whereas the curve slope gets flatter with IgG density, and the overall amount of prey associated with phagocytes is only significantly increased with the IgG density. The middle row explores at what MOPs there are 10, 50 or 90% persistently associated phagocytes. Here, the trend looks similar at the lower association scenarios, with expectedly increased IgG density leading to lower MOPs required for reaching a given persistent association. At 90% association, the IgG density appears to have little effect on the outcome. In the lower panel, we analyzed the number of PxP at MOP_10_,MOP_50_, and, MOP_90_, which showed similar levels for each degree of association. There are small differences within each group, but in general the number of prey per phagocyte seems more dependent on association than on IgG density. Overall the association curve characteristics indicate that the primary effect of increased IgG density lies in a higher persistent association, and not primarily affecting the fate of prey that has been associated with phagocytes.

To further explore the effects of prey IgG density, we analyzed persistent association and the fate of associated prey (**Fig. 4E**). First, we looked at the probability to interact with at least one prey, phagocytic interaction, which exponentially increases with IgG density (R^2^=0.98). That means that to evoke 50% of maximal persistent association only one third of the number of beads was needed when opsonized with 1000 µg/ml compared to 1 µg/ml. However, the average number of associated beads per interacting phagocyte was 2-3 beads independent of IgG density (**Fig. 4E**). Still, internalization appears to increase exponentially as well, but not as strongly as phagocytic association (R^2^=0.68). From 1 to 1000 µg/ml the average number of beads internalized increased two-fold. As expected, adhesion had an inverted relationship compared to internalization. Overall, our findings indicate that the association capacity of a phagocyte population increases exponentially with IgG density combined with a smaller, but increased internalization capacity of prey-associated phagocytes.

### Association characteristics across prey reveal biological differences

A conclusion from this study is that it normally would be very difficult to compare phagocytosis across different types of prey, including species and strains if they are expected to have an effect on phagocyte association. However, by normalizing for persistent association using the PAN-method, it would be possible to actually compare quite different types of prey in a quantitative manner. Here we compared the gram-positive cluster-forming cocci *S. aureus*, with the gram-negative bacilli *E. coli*, and streptavidin-coated beads, all opsonized with 10 mg/ml IgG.

Overall, phagocytosis appears to be different when changing the prey (for flow cytometry scatter plots see **Supp. Fig. 3**). The persistent association curves either have similar slopes but different top-levels, or a different slope compared to the others (**Fig. 5A**). The number of associated prey is also different, with the beads at a much lower level than the others (**Fig. 5B**), even when normalized for potential differences in fluorescence signal between prey the top right figure in **5C** clearly shows that the overall amount of prey associated with phagocytes is very low for the beads compared to bacteria. This is also consistent with that much more beads are required to reach a similar level of persistent association, whereas the bacteria behave similarly in terms of MOP for a given association (**Fig. 5C**). The differences in surface properties between the prey might be a natural reason why beads are not as good phagocytic target as bacteria. However, interestingly enough beads have the same maximum association value as *E. coli*, around 100% of the phagocytes will associate with them at high levels of MOPs, while *S. aureus* is saturated around 80%. Looking at prey per phagocyte reveals that *E. coli* are associating better at lower levels of interaction (MOP_10_ and MOP_50_), whereas *S. aureus* are more prone to be associated at MOP_90_ than the other prey. At this point, we do not know what the differences in PA_max_, Hill coefficient or association characteristics mean mechanistically, but we know that these differences are indicative of a biological difference in terms of phagocytosis. This shows that the sensitivity and comparative quantification that PAN-based analysis offers, opens up for further mechanistic studies or careful quantitative comparisons that might explain inherent differences in phagocytosis across prey.

**Figure 5.**
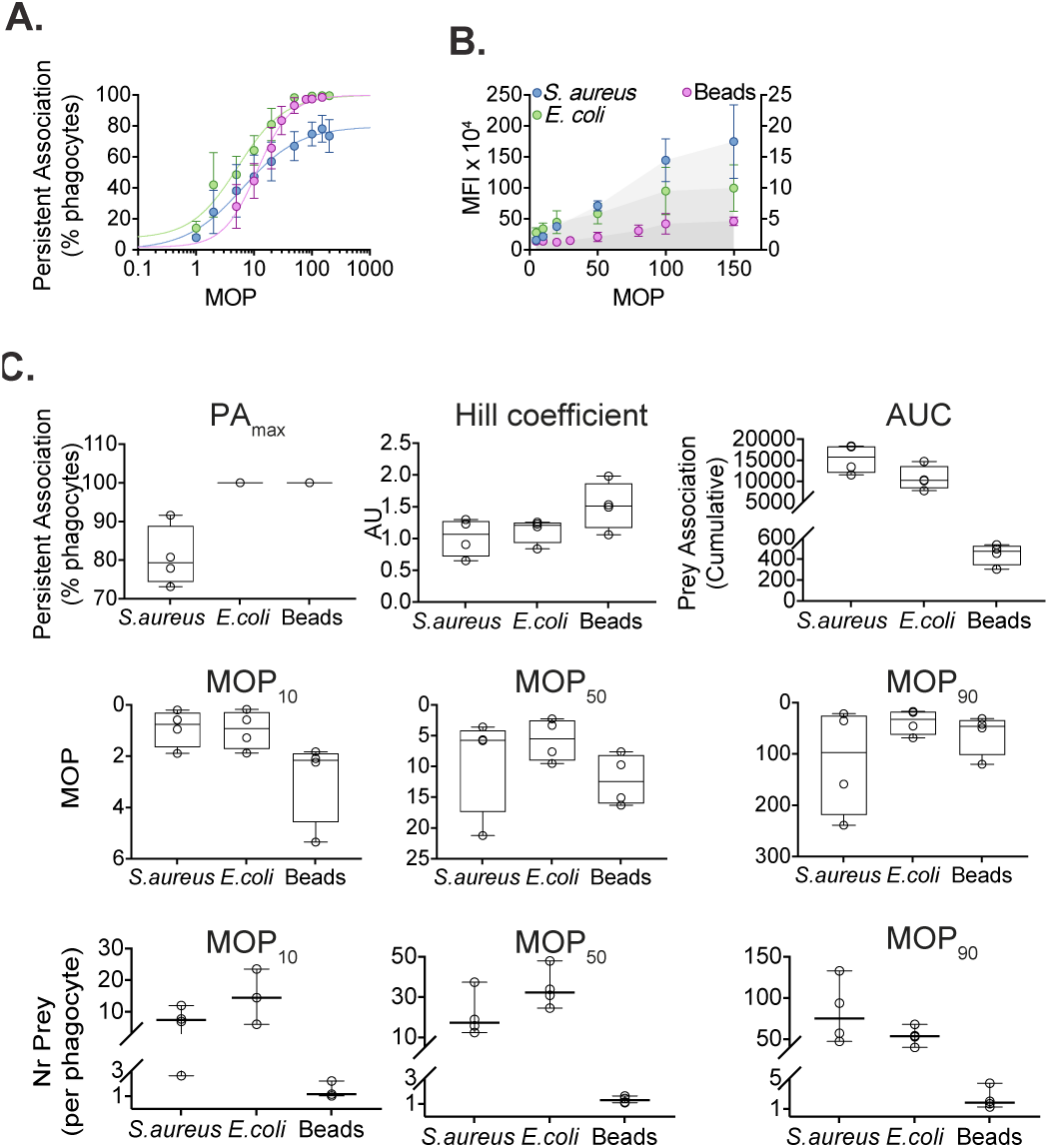
Association characteristics across prey reveal biological differences. (A-C) *S. aureus, E. coli*, and streptavidin-coated beads all opsonized with 10 mg/ml IgG were incubated with THP-1 cells for 30 min with MOP 0-200 for the bacteria and 0-150 for the beads prior to paraformaldehyde fixation. Single preys were run separately allowing for conversion of the fluorescence signal to a number of preys. Data were acquired through flow cytometry and are represented as mean ± SD, n=4. (A) The average of the persistent association curves for each prey (R^2^ mean ± SD, n=4: *S. aureus* 0.96 ± 0.03, *E. coli* 0.94 ± 0.09, beads 0.98 ± 0.03) (B) Corresponding Median Fluorescence Intensity (MFI) to association curves in A. Left Y axis is showing *S. aureus* and *E. coli* while the right Y axis represents bead MFI. (C) The characteristics of the dose-response curves for each prey as retrieved using the analysis template provided (Supp. Fig 4). For area under curve (AUC) quantification, the MFI values were normalized for each prey and calculated between the same range of MOP (0-150) to be comparable. All data points are shown, the distribution is visualized with box plots with whiskers ranging from minimum to maximum and a line marking the median, n=4. The last row shows all data points, their median with 95 % confidence interval, n=4 except at MOP_10_ where n=3 for E. coli and beads.

## Discussion

Phagocytosis is measured as a functional outcome in many fields (**Supp. Fig. 1**), and could even be considered a standard assay for many research areas. Despite this, there is no gold standard and no established way of doing a robust comparison across experiments or laboratories. In the present work, we have provided a new framework for the measurement of phagocytosis, including theory, terminology, as well as analysis guidelines and templates (**Supp. Fig. 4**). A major benefit to the method we introduce here is increased robustness and sensitivity, and also the fact that it can be readily implemented into most existing methods to measure phagocytosis. Our hope is that this will allow for both better data when measuring phagocytosis as well as provide the option for cross-experiment and cross-laboratory standardization.

The PAN method is fundamentally based on dose-response curve analysis, commonly applied in a number of fields, and particularly in pharmacology (Holford and Sheiner, 1981). There it has become an essential way of doing analysis, with effective concentration (EC_50_) or inhibitory concentration (IC_50_) numbers commonly used to evaluate the pharmacological effect and has provided an analytical tool to compare cross-laboratory, across different compounds and over time. The MOP_50_ used in PAN could potentially be used in a similar fashion. Additionally, translating individual data points into a mathematical function also comes with a number of benefits, where curve characteristics can be quantified and compared. MOP_10_, MOP_50_ and MOP_90_ allows for the exploration of scenarios where phagocytes are encountering various amounts of prey, and what the physiological effect could be. The slope or Hill coefficient offers both intriguing ramifications, where a coefficient different from 1 indicates either positive or negative feedback (Hill, 1910), and is important to quantify as shown in IC_50_ studies on HIV, where effect by resistance mutations have been overlooked due to lack of slope analysis (Sampah et al., 2011). To our knowledge the impact of how interacting with one prey affects subsequent interactions with additional prey, has not been studied for phagocytosis; Hill coefficient analysis would offer a guide of where to start to look for such studies. The PAN method is especially valuable when looking at different bacterial mutants or strains that would affect surface properties (Happonen et al., 2019), and thus potentially directly affect persistent association. A key aspect that should be emphasized is that the MOP curves not only serve as an analysis in itself, but can also provide a guide to the dynamic range of the system, and at which MOP more detailed analysis, such as microscopy, should be performed, to improve sensitivity and potentially save unnecessary experimental set up time.

A comparative meta-analysis is feasible with a standardized approach. That way it would be possible to create a database with phagocytosis characteristics for phagocytes and prey from many different sources. This could, for instance, be PxP and MOP_50_ numbers for different phagocytes and strains of bacteria under various opsonizing conditions. Such a database could then be used for the mining of phagocytosis data to identify important pre-clinical as well as clinical data. Systems with limited, sensitive data and potentially high heterogeneity such as different clinical isolates and patient material would benefit not only from increased assay sensitivity, but also from generating comparable results stable over time that could be preserved in such a database. In the infection medicine field, pre-clinical and clinical researchers come together in the important undertaking to develop alternative therapeutics to antibiotics. Identifying functional monoclonal antibodies that can be used to treat infections would be an obvious use for the PAN-method.

In summary, we have established a universal analytical method that can be used across different systems to normalize for factors affecting the association between phagocyte and prey, where adhesion and internalization can be analyzed separately with improved robustness and assay sensitivity.

## Materials and Methods

### Microbe strains

*Staphylococcus aureus* Cowan-1 and *Escheria coli* DH5alpha (Invitrogen) were cultured under shaking conditions at 37°C with 5% CO_2_ atmosphere in THY-medium (Todd Hewitt Broth; Bacto; BD, complemented with 0.2% (w/v) yeast) or LB-medium (Luria-Bertani;Sigma-Aldrich), respectively. The bacteria were harvested at optical density OD_620_ 0.4.

### Cell lines

THP-1 cells (Tsuchiya et al., 1980) (ATCC TIB-202, male) were maintained in Roswell Park Memorial Institute medium 1640 (Sigma-Aldrich) supplemented with 10% fetal bovine serum (Gibco), 1% Penicillin-Streptomycin (ThermoFisher) and 2 mM GlutaMAX (Life Technologies). Cells were grown at 37°C in a 5% CO_2_ atmosphere. The cell density was kept between 0.2 -1.0 × 10^6^ cells/ml with a viability over 95% and were kept up to 3 months before thawing a new aliquote of fresh cells.

### Beads

Flash Red-labeled 1 µm microspheres coated with streptavidin (Polysciences Europe Bangs Laboratory. Inc, catalog nummer CFFR004) were washed 3 times in 1 ml Na-medium (5.6 mM glucose, 127 mM NaCl, 10.8 mM KCl, 2.4 mM KH_2_PO_4_, 1.6 mM MgSO_4_, 10 mM HEPES, 1.8 mM CaCl_2_; pH adjusted to 7.3 with NaOH) at 20 000 x *g* for 2 min and kept protected from light at 8°C prior to opsonization. All chemicals were from Sigma-Aldrich.

### Heat-killing of bacteria

After harvesting the bacteria, they were washed 3 times with 1 ml Na-medium at 2000 x *g* for 2-5 min. Heat-killing were induced at 80°C for 5 min in a heat-block, vigorously shaking, followed by cooling the bacteria rapidly on ice. Bacteria not used directly were stored in fridge up to two weeks.

### Labeling of bacteria

Heat-killed bacteria were stained at 37°C with gentle shaking and protected from light for 30 min either with 2 µg/ml DyLight™650 (ThermoFisher) or 5 μM Oregon Green 488-X succinimidyl ester (Invitrogen), depending on if fixation of phagocytosis was planned, if so the former staining was used. The samples were washed once with Na-medium (1 min, 5000 x *g*, swing-out rotor) and if needed, sonicated for up to 4 min (VialTweeter; Hielscher) to disperse any large aggregates of bacteria which was confirmed by microscopy. The samples were then gently spun down (200 x *g*, 2 min, swing-out) and continued with the supernatant to remove any remaining bacterial aggregates.

### Opsonization of prey

Intravenous immunoglobulin (IVIG; Octagam, Octapharma) was centrifuged 2 min at ∼20 000 x *g* to precipitate any aggregates. The prey were opsonized with 0-10 mg/ ml of IVIG at 37°C with gentle shaking and protected from light for 30 min. Any unbound IVIG was washed away with 1 ml Na-medium five times (1 min, 5000 x *g* for bacteria and 20 000 x *g* for beads, swing-out) for all experiments except when evaluating the effect IgG density has on phagocytosis.

### Preparation of THP-1 cells

On the day of experiments, the cell density was kept between 0.50-0.7×10^6^ cells/ml. Medium was changed to Na-medium (5 min, 500 x *g* fixed rotor) and cells were kept on ice until start of the phagocytosis assay. In experiments without fixation, dead cells were labeled with 2 µM of the impermeable cell membrane dye DRAQ7™ (Abcam). In experiments with fixation, cells were instead labeled according to manufacturer’s protocol with LIVE/DEAD™ Fixable Violet Dead Cell Stain Kit, for 405 nm excitation (ThermoFisher) except when evaluating the effect IgG density has on phagocytosis.

### Phagocytosis assay

The concentration of prey and THP-1 cells were measured prior to phagocytosis by flow cytometry (Accuri C6, BD or CytoFLEX, Beckman-Coulter), if needed, prey were sonicated in advance. The concentration of THP-1 cells was set for each experiment and ranged between 1×10^5^-2×10^6^ cells per sample. The cells were added on ice to their prepared samples, at least 7 different MOP with the range 0–200 were prepared for each experiment. Incubation was performed in Na-medium at 37°C on a heating-block with moderate shaking protected from light. Phagocytosis was haltered by either putting the samples on ice, adding of 1 µM Cytochalasin D (ThermoFisher) for 15 min or by fixing them in 1% PFA (ThermoFisher) for at least 30 min.

In general, the incubation time and volume was 30 min respectively 150 µl if not specified otherwise. Phagocytes and prey were centrifuged (3000 x *g*, 1 min, including 30 sec of acceleration) together prior to incubation when evaluating the effect of centrifugation on association. When possible, 96-well-plates, 8-channel multi-pipet and low-binding pipet tips and Eppendorf tubes were used.

### Staining and fixation

Fixation was performed in 1% paraformaldehyde (PFA; ThermoFisher) for at least 30 min on ice protected from light. Post-fixation samples were incubated with 50 mM glycine and 5% BSA for at least 10 min at room temperature, except when evaluating the effect IgG density has on phagocytosis where only glycine was used.

THP-1 cells were labeled with mouse antibody anti-human CD18 (PE BioLegend or BV421 BD) (1 µl per 200 000 cells) for at least 10 min at room temperature. The extracellular bacteria were labeled with 1:1000 Fab specific DyLight 488-conjugated AffiniPure F(ab’) Fragment Goat Anti-human IgG (Jackson ImmunoResearch). Fluorescent Atto 488nm-biotin (0.008 µg/sample, Sigma Aldrich) was used to label extracellular beads.

### Data acquisition

Flow cytometric acquisition was performed using three different flow cytometers. Accuri C6 (BD) equipped with 640 nm and 488 nm lasers, CytoFLEX (Beckman-Coulter) with 405 nm, 488 nm, and 638 nm with 450/45 PB450, 525/40 KO25, respectively 525/40 FITC, 585/42 PE, and 660/10 APC, 780/60 APC-A750 filters and FACSVerse (BD) with 640 nm and 488 nm lasers with filters 527/32 FITC, 586/42 PE, 660/10 APC and 783/56 APC-Cy7. For each experiment threshold, gain and velocity was set. Acquisition was set to analyze at least 5 000 events of the target population.

### Data analysis

Flow cytometry data were analyzed using FlowJo version 10.2 (Tree Star). Compensation was performed using in-built matrices in the CytoFLEX or FACSVerse when needed. The general gating strategy was similar for all experiments (see **Supp. Fig. 5-7**). Dead cells were excluded by being extremely positive (a population at least 2 logs higher than all other signals) for DRAQ7™ or for the LIVE/DEAD violet signal. Doublets were excluded by gating on FSC-H versus FSC-A. THP-1 cells were gated on forward and side scatter and CD18 when used. We defined association as a THP-1 cell being positive fluorescence corresponding to a prey, Oregon Green, DyLight 650 or Flash Red.

### PAN-method

#### MOP_50_-curve

Association, the percentage of phagocytes positive for at least one prey, was plotted versus MOP resulting in curves that was analyzed using Prism 8.2.1 (GraphPad) inbuilt non-linear regression analysis tool named “Agonist vs. response --Variable slope (four parameters) or Find ECanything”. Least squares regression was used with no special handling for outliers and no weighting. Parameters of the curves were constrained as follow: bottom – more than 0, to between 0-100 and EC50 had to be bigger than 0. If technical replicates were used the mean was used to generate one curve, for each replicate one curve was generated but the experiment is visualized with one average for the whole experiment. The characteristics of the curves are generated when the curve is created except for Area Under Curve (AUC) which is based on the prey MFI for the whole MOP range. When comparing between different preys or labeling techniques the signal was first normalized by dividing it by the signal for a single prey and then compared at same MOP range.

#### Normalization

The fluorescent intensity for association and adhesion was plotted separately versus MOP expressed in mean, median or mode. Normalization was performed by interpolating the MFI corresponding to the MOP which generated by above mentioned analysis evoking half of the maximal response (MOP_50_). Interpolation was done depending on the best fit either by non-linear regression or linear-regression between the two MOPs that were in-between MOP_50_ or when needed with a manual read out.

#### Internalization and PxP

The fluorescent signal of a prey unit was determined by measuring the single prey unit by flow cytometry. This could then be used to determine the ratio between the fluorescent staining thus converting adhesion signal to same unit as association signal by multiplying it with the ratio allowing us to estimate the internalized signal through subtraction. Thus, the total prey signal subtracted by the adhered prey signal results in internalized prey signal. The number of prey per phagocyte, PxP, could be quantified by dividing each signal with the corresponding prey unit signal. Here, assuming one prey unit is one prey.

### Statistical analysis

The statistical details of experiments can be found in each figure legend, but mean and SD is used if not otherwise specified. All statistical analysis performed in this paper was done using Prism 8.2.1 (GraphPad). Correlation was evaluated using non-parametric Spearman, two tailed with alpha 0.05 and non-linear regression was performed using semi-log fit with the mean of the y-replicates. The signal-to-noise (SNR) ratio was quantified for each level by dividing each different IgG-density signal with the corresponding background, here the non-opsonized samples were set as background.

### Additional Resources

Github page with templates, see link.

## Acknowledgments

We acknowledge the Oonagh Shannon Lab for help and access to their flow cytometer. PN and TdN was funded by the Royal Physiographic Society. MS and PN was funded by the Gyllenstierna-Krapperup Foundation. PN was funded by the Swedish Research Council (VR), Swedish Society of Medicine (SLS), the Crafoord Foundation, the Schyberg Foundation, the Groschinsky Foundation, and the Österlund Foundation.

