## Supplementary figures for "High-sensitivity assessment of phagocytosis by persistent association-based normalization"

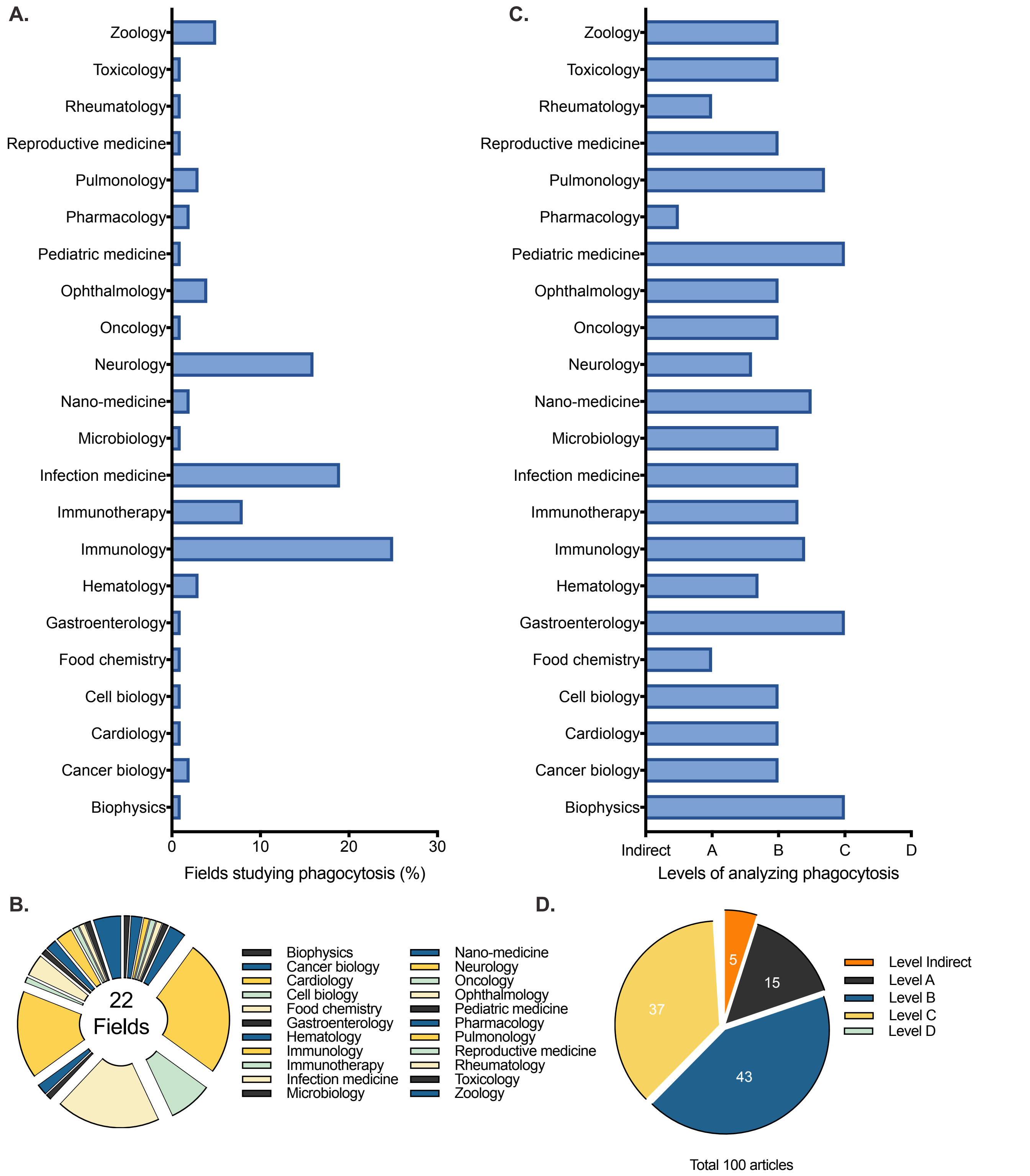


**Supplementary figure 1. Assessments of phagocytosis in the literature.**
A literature search in the years of 2013-2018 on 100 papers having a phagocytosis assay as part of their method resulted in papers from at least 22 different fields. (A-B) The distribution of the different fields visualized as diagram (A) and a pie-chart (B). (C) The average level of each fields' phagocytosis assessment. (D) The distribution of the different levels of phagocytosis assessment visualized as a pie-chart. Different types of indirect methods to study phagocytosis belongs to Level Indirect, such as plate-based killing assays. Level A is assessing the whole phagocytic population while Level B can distinguish between associating and non-associating phagocytes, Level C further separates between adhesion and internalization. Level D is introduced here (PAN method) by determining MOP50 before quantifying phagocytosis.

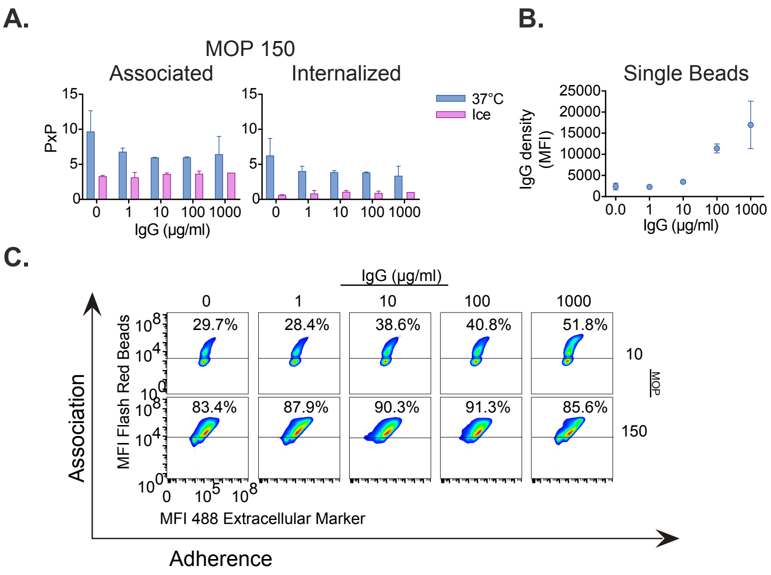


**Supplementary figure 2. Relationship between IgG density and concentration, temperature and phagocytosis.**

(A)As a control experiment THP-1 cells were incubated (30 min, 150 µl, MOP 150, 37°C or on ice) with 1 µm Far Red streptavidin beads opsonized with 0-1000 µg/ml of human polyclonal IgG. Alexa488-conjugated biotin to mark extracellular beads post fixation. Single beads were run separately to allow conversion of the fluorescence signal to number of beads, prey per phagocyte (PxP). Data were acquired through flow cytometry and are presented as mean ± SD n=2 (1000 µg/ml ice n=1). Compared to 37°C, phagocytosis on ice clearly decreases internalization and to some degree also association. (B) Single beads opsonized with 0-1000 µg/ml of human polyclonal IgG were run separately to confirm increasing IgG density on the bead surface corresponding to increased opsonin concentration. This was assessed by incubating opsonized beads with DyLight 488-conjugated f(ab')2 fragment anti-human IgG+IgM for 30 min 37°C (1:400). Data were acquired through flow cytometry and are presented as mean ± SD n=4. (C)Representative flow cytometry scatter plots of the experiment evaluating phagocytosis at different IgG concentrations (Figure 3 and 4) at MOP 10 and 150. Median fluorescence intensity (MFI) on the y-axis, Far Red, corresponds to the signal from all associated beads and X-axis signal corresponds to adhered beads labeled with Alexa488-conjugated biotin.

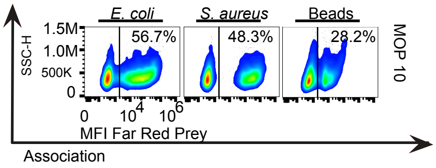


**Supplementary figure 3. Phagocytosis of different prey.**

(A) Representative flow cytometry scatter plots of the experiment evaluating phagocytosis with different preys, *S. aureus, E. coli* and streptavidin-coated beads all opsonized with 10 mg/ml IgG at MOP 10 (Figure 5). Median fluorescence intensity (MFI) on the X-axis, Far Red, corresponds to the signal from all associated prey

**A.**

Templat_prism_advance monday 28 october 2019 09:54

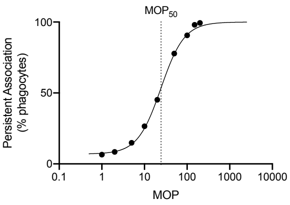


Curve
$R^2$: 0.9983
Top: 100.0
Bottom: 6.953
Hillslope: 1.585
$MOP_{50}$: 24.45

At $MOP_{50}$
PxP associated: 27.226
PxP adhered: 21.781
PxP internalized: 5.445

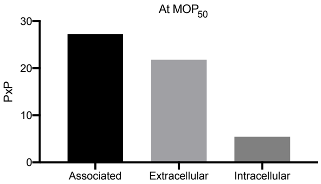


| | Curve | MFI at $MOP_x$ | PxP at $MOP_x$ | AUC MFI x MOP | AUC PxP x MOP |
|---|---|---|---|---|---|
| ECF | 24.4500 | | | | |
| F | 50.0000 | | | | |
| HillSlope | 1.5850 | | | | |
| Bottom | 6.9530 | | | | |
| Top | 100.0000 | | | | |
| R squared | 0.9983 | | | | |
| Associated Prey | | 21780.79333333330 | 27.22599166666670 | 8463874.00 | 10580.00 |
| Extracellular Prey | | 10890.39666666670 | 21.78079333333330 | 4231937.00 | 8464.00 |
| Intracellular Prey | | 5445.19833333333 | 5.44519833333333 | 2115968.00 | 2116.00 |

| MOP | % Association |
|---|---|
| 1 | 6.57 |
| 2 | 8.47 |
| 5 | 14.90 |
| 10 | 26.50 |
| 20 | 45.20 |
| 50 | 77.80 |
| 100 | 90.60 |
| 150 | 98.00 |
| 200 | 99.40 |

| MOP | MFI Associated Prey | MFI Extracellular Prey | MFI Intracellular Prey |
|---|---|---|---|
| 1.00 | 13193.00 | 6596.50 | 3298.25 |
| 2.00 | 16033.00 | 8016.50 | 4008.25 |
| 5.00 | 15024.00 | 7512.00 | 3756.00 |
| 10.00 | 20792.00 | 10396.00 | 5198.00 |
| 20.00 | 19484.00 | 9742.00 | 4871.00 |
| 50.00 | 34968.00 | 17484.00 | 8742.00 |
| 100.00 | 39821.00 | 19910.50 | 9955.25 |
| 150.00 | 55109.00 | 27554.50 | 13777.25 |
| 200.00 | 66971.00 | 33485.50 | 16742.75 |

| Constant | Value |
|---|---|
| Prey Signal Association | 800 |
| Prey Signal Extracellular | 500 |
| Prey Signal Intracellular | 1000 |

Analyzed with PAN, method created by Therese de Neergaard & Pontus Nordenfelt 2019

**B.**

Templat_prism_basic wednesday 23 october 2019 15:20

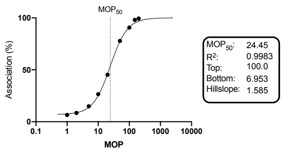


| | Curve | MFI |
|---|---|---|
| Best-fit values | | |
| Bottom | 6.9530 | |
| Hillslope | 1.5850 | |
| Top | 100.0000 | |
| EC50 | 24.4500 | 21780.7933333333 |
| R squared | 0.9983 | |
| AUC | | 8463874.0000000000 |

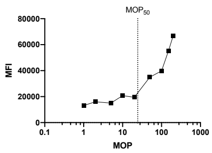


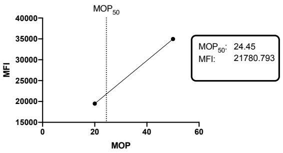


| MOP | % Association |
|---|---|
| 1 | 6.57 |
| 2 | 8.47 |
| 5 | 14.90 |
| 10 | 26.50 |
| 20 | 45.20 |
| 50 | 77.80 |
| 100 | 90.60 |
| 150 | 98.00 |
| 200 | 99.40 |

| MOP | MFI |
|---|---|
| 1.00 | 13193.00 |
| 2.00 | 16033.00 |
| 5.00 | 15024.00 |
| 10.00 | 20792.00 |
| 20.00 | 19484.00 |
| 50.00 | 34968.00 |
| 100.00 | 39821.00 |
| 150.00 | 55109.00 |
| 200.00 | 66971.00 |

Analyzed with PAN, method created by Therese de Neergaard & Pontus Nordenfelt 2019

**Supplementary figure 4. Examples of PAN templates.**

Illustrations of the output from PAN templates in Prism 8 (Graph Pad) provided by the authors, in the form of basic (B) and advanced (A) templates. Both templates determine the persistent association curve, its characteristics and interpolate data at MOP50. In addition, the advanced template allows for assessment at MOPx of choice and to convert data into prey per phagocyte PxP.

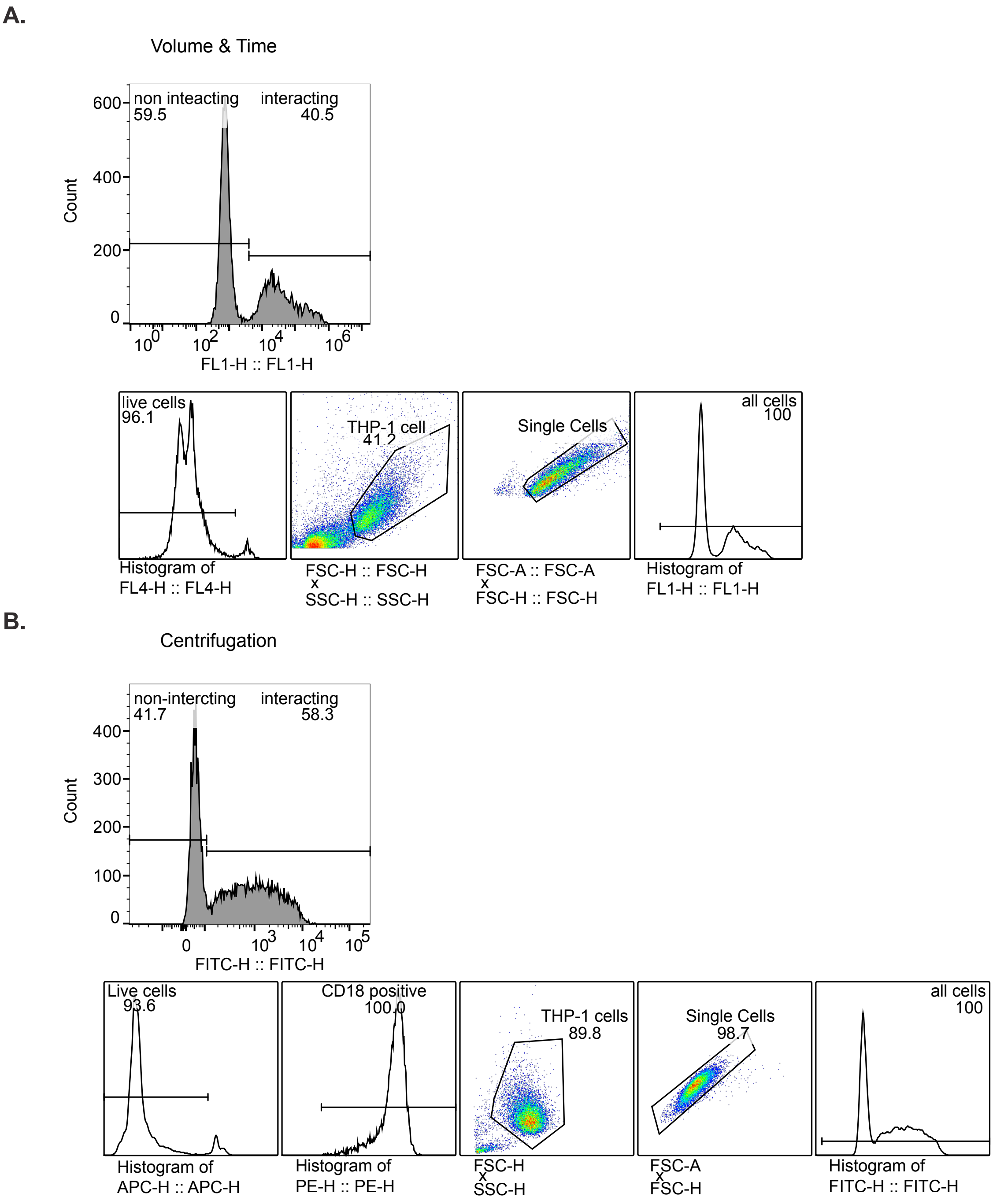


**Supplementary figure 5. Gating strategy for phagocytosis at different experimental parameters.**
The gating strategy for the experiments in figure 1 and 2, where phagocytosis of *S. aureus* with different volumes, time-points and centrifugation were evaluated. Volume and time were gated exactly the same while centrifugation has an additional gating for CD18 positivity after live cell gating. Draq 7, a non-cell-membrane permeable Far-Red fluorescent dye was used to exclude dead THP-1 cells. Followed by gating on positivity for anti-CD18-PE for centrifugation experiment or directly gated on forward and side scatter, FSC-H vs SSC-H. Doublets were excluded through a height versus area gating on FSC. Subsequently, a gate for all cells was created including all data above zero fluorescence. Persistent association was then determined by the percentage of all cells positive for Oregon Green labeled preys, FL1-H.

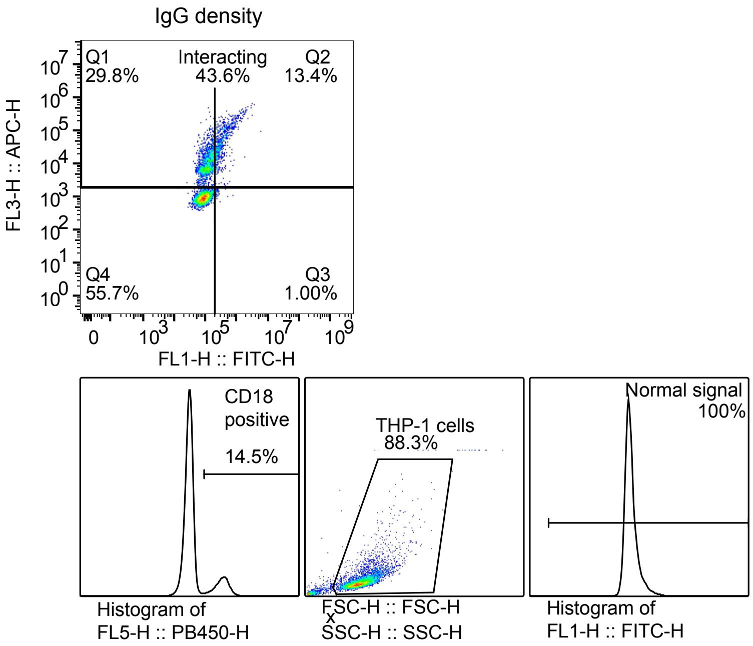


**Supplementary figure 6. Gating strategy for phagocytosis of beads opsonized with different IgG density.**
The gating strategy for the experiments in figure 3 and 4, where phagocytosis with different IgG density was evaluated differs slightly from previous stated gating strategy. First anti-CD18-BV421 positive cells were gated on pacific blue, FL5-H, followed by gating on FSC-H vs SSC-H, before a gate for all cells was created including all data above zero fluorescence. Persistent association was then determined by the percentage of all cells positive for Far Red positive streptavidin-coated beads, FL3-H, and extracellular beads positive for 488-conjugated biotin, FL1-H.

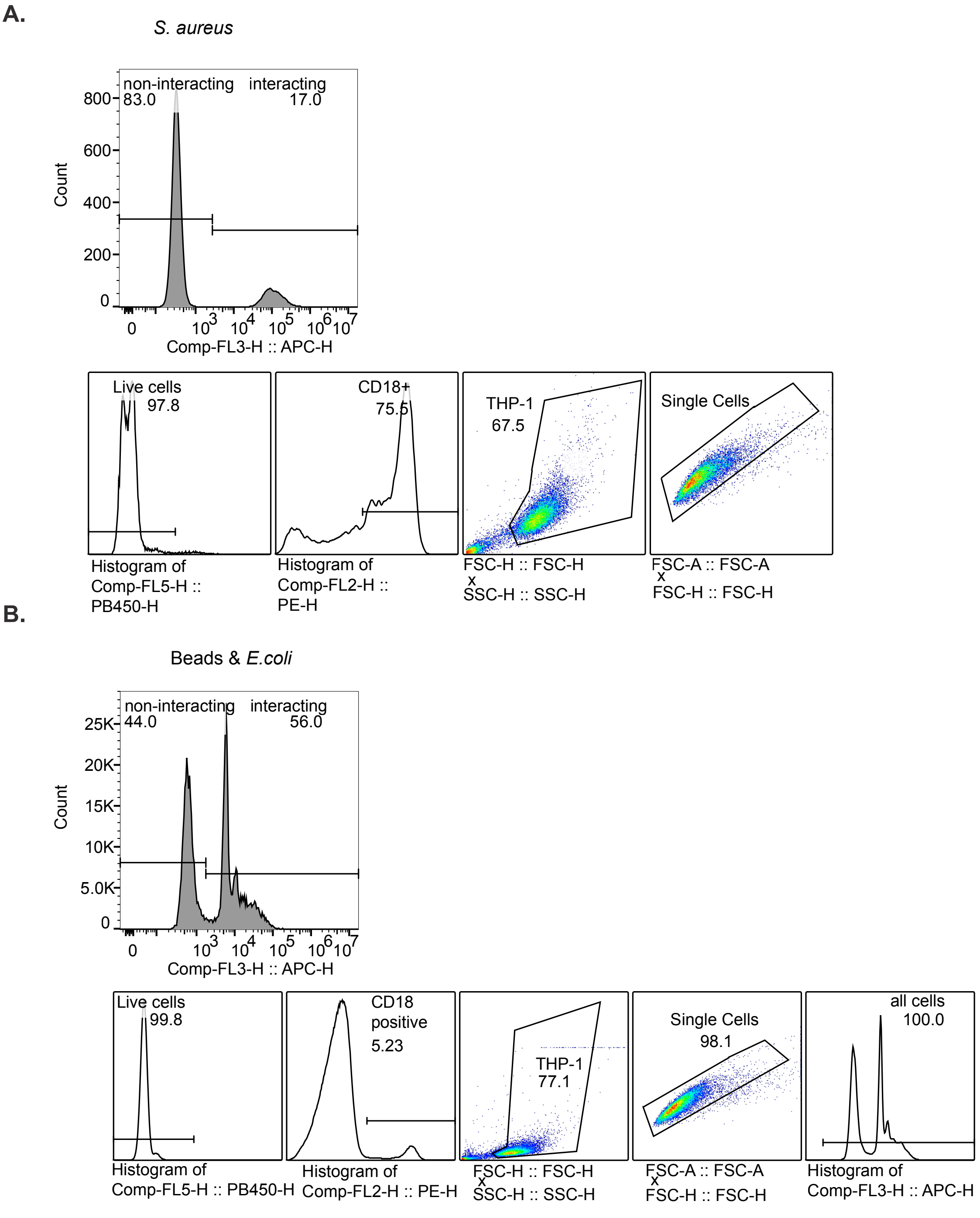


**Supplementary figure 7. Gating strategy for phagocytosis of different prey.**
The gating strategy for the experiments in figure 5, where phagocytosis with different preys opsonized with 10 mg/ml of IgG were compared. Streptavidin-coated beads and *E. coli* gating strategy (B) was the same, while *S. aureus* (A) went directly from excluding doublets to determine the persistent association, beads and E. coli first had a gate on all cells with fluorescence above zero. For all preys, dead cells were excluded by being positive in the violet channel, FL5. The gating was followed by selecting cells being positive for anti-CD18-PE, FL2. Then FSC-H x SSC-H was gated on before excluding doublets by gating on area versus height, FSC. Persistent association was determined by the percentage of all cells positive for prey in Far Red channel, FL3.
